# Diversity, composition, altitude, and seasonality of high-altitude windborne migrating mosquitoes in the Sahel: Implications for disease transmission

**DOI:** 10.1101/2022.08.19.504620

**Authors:** AS Yaro, Y-M Linton, A Dao, M Diallo, ZL Sanogo, D Samake, Y Ousman, C Kouam, BJ Krajacich, R Faiman, R Bamou, J Woo, JW Chapman, DR Reynolds, T Lehmann

**Author notes:** **Correspondence: Tovi Lehmann**.

## Abstract

Recent studies have reported *Anopheles* mosquitoes captured at high-altitude (40–290 m above ground) in the Sahel. Here, we describe this migration modality across genera and species of African Culicidae and examine its implications for disease transmission and control. As well as *Anopheles*, six other genera—*Culex, Aedes, Mansonia, Mimomyia, Lutzia*, and *Eretmapodites* comprised 90% of the 2,340 mosquitoes captured at altitude. Of the 50 molecularly confirmed species (N=2,107), 33 species represented by multiple specimens were conservatively considered high-altitude windborne migrants, suggesting it is a common migration modality in mosquitoes (31–47% of the known species in Mali), and especially in *Culex* (45–59%). Overall species abundance varied between 2 and 710 specimens/species (in *Ae. vittatus* and *Cx. perexiguus*, respectively). At altitude, females outnumbered males 6:1, and 93% of the females have taken at least one blood meal on a vertebrate host prior to their departure. Most taxa were more common at higher sampling altitudes, indicating that total abundance and diversity are underestimated. High-altitude flight activity was concentrated between June and November coinciding with availability of surface waters and peak disease transmission by mosquitoes. These hallmarks of windborne mosquito migration bolster their role as carriers of mosquito-borne pathogens (MBPs). Screening 921 mosquitoes using pan-*Plasmodium* assays revealed that thoracic infection rate in these high-altitude migrants was 2.4%, providing a proof of concept that vertebrate pathogens are transported by windborne mosquitoes at altitude. Fourteen of the 33 windborne mosquito species had been reported as vectors to 25 MBPs in West Africa, which represent 32% of the MBPs known in that region and include those that inflict the heaviest burden on human and animal health, such as malaria, yellow fever, dengue, and Rift Valley fever. We highlight five arboviruses that are most likely affected by windborne mosquitoes in West Africa: Rift Valley fever, O’nyong’nyong, Ngari, Pangola, and Ndumu. We conclude that the study of windborne spread of diseases by migrating insects and the development of surveillance to map the sources, routes, and destinations of vectors and pathogens is key to understand, predict, and mitigate existing and new threats of public health.

## Introduction

High-altitude windborne migration of large insects such as locusts has been recognized over millennia, however, over the past decades, our knowledge of this behavior has greatly expanded thanks to systematic studies using aerial sampling, radar, and other methodologies (Reynolds & Riley, 1988; Chapman *et al.*, 2011, 2015). Radar studies have been key to estimate direction, speed, altitude, and magnitude of migrating insects, but they seldom provide species-level information (Drake & Reynolds, 2012). Given the enormous number of insects involved, the distances they cover, and their interactions with other organisms, windborne migration influences food security (Riley *et al.*, 1987; Chapman *et al.*, 2002, 2004a, 2015; Maiga *et al.*, 2008), public health (Garrett-Jones, 1950; Garms *et al.*, 1979; Kay & Farrow, 2000; Johansen *et al.*, 2003; Reynolds *et al.*, 2006; Huestis *et al.*, 2019), and ecosystem vigor (Hu *et al.*, 2016; Wotton *et al.*, 2019). Although windborne flight at altitude is common in many groups of insects including Diptera (Garms *et al.*, 1979; Chapman *et al.*, 2004b; Sanders *et al.*, 2011; Eagles *et al.*, 2013; Miao *et al.*, 2013; Wotton *et al.*, 2019), most studies on mosquito movements have been focused on host location and on movement at a village scale (Gillies, 1961; Costantini *et al.*, 1996; Toure *et al.*, 1998; Takken & Knols, 1999) under the premise that dispersal of most mosquito species does not exceed 3–5 km, and long-range movements represent rare ‘accidental events’ that are of minimal epidemiological importance (Service, 1997; Harrington *et al.*, 2005; Verdonschot & Besse-Lototskaya, 2014). The term dispersal is commonly used in mosquito literature (Service, 1997), but the definition of migration as ‘persistent movements unaffected by immediate cues for food, reproduction, or shelter, with a high probability of relocating the animal in a new environment’ (Dingle & Drake, 2007; Chapman *et al.*, 2015) is apt here and will be used hereafter.

Recent studies in the Sahel revealed that *Anopheles* mosquitoes, like other windborne insect migrants, regularly engage in high-altitude windborne migration: e.g., 40–290 m above ground level (agl), between July and November (Huestis *et al.*, 2019; Faiman *et al.*, 2020; Florio *et al.*, 2020). Furthermore, gravid females (i.e., females with mature eggs, following a blood meal) predominated among mosquito migrants. The number of individuals that cross a 100 km line perpendicular to the wind at altitude was found to range between tens of thousands to millions per year, depending on mosquito species. Finally, aided by winds at altitude, these mosquitoes cover tens to hundreds of kilometers per night (assuming up to nine-hour nightly flights). These findings pertain to Sahelian *Anopheles* species and were interpreted in the context of malaria transmission and control. Here, we expand the scope of our first analysis (Huestis *et al.*, 2019) and include the full collection of mosquitoes that were intercepted at altitude over the Sahel to better assess the implications of this migration modality in mosquitoes to public health and ecosystem stability in West Africa. Specifically, we identify the species intercepted at altitude, and for each species estimate their sex ratio and gonotrophic state composition, mean flight altitude, and seasonality – all parameters that impinge on risk of infection with mosquito-borne pathogens (MBPs)(Garrett-Jones & Shidrawi, 1969). We report on the initial screening of these mosquitoes to infection with species of *Plasmodium*, which infect a range of vertebrate hosts. Focusing on the windborne mosquito species that have been (previously) reported as vectors of MBPs in West Africa, we rank both the risk of the pathogens for windborne spread and the mosquito species in terms of their total contributions to overall MBP spread. Our results indicate that windborne migration of mosquitoes is a key driver shaping the epidemiology of many mosquito-borne diseases and that this newly recognized aspect of the vector and MBP ecology should be considered in predicting disease spread as well as in planning disease control strategies.

## Materials and Methods

The study area has been described in detail previously (Lehmann *et al.*, 2010, 2017; Huestis *et al.*, 2012; Dao *et al.*, 2014), as has the field and most laboratory methods used in this study (Huestis *et al.*, 2019; Florio *et al.*, 2020; Sanogo *et al.*, 2021). Below is a brief description of the field and laboratory operations previously reported and a detailed description of other procedures. Aerial sampling stations were placed in four Sahelian villages with traditional mud-brick houses, surrounded by fields, set in a dry savanna, consisting of grasses, shrubs, and scattered trees: Thierola (13.6586, – 7.2147) and Siguima (14.1676, – 7.2279) were sampled between March 2013 to November 2015; Markabougou (13.9144, – 6.3438) sampled between June 2013 to June 2015, and Dallowere (13.6158, – 7.0369) sampled between July to November 2015. Sampling was not carried out in January and February. Over 90% of the rains fall in the wet season (June—October, ~ 550 mm annually), forming temporary puddles and pools that usually dry by November. Rainfall during the dry season (December—May) is negligible (0—30 mm)(Touré *et al.*, 2017).

Aerial insect sampling was conducted using sticky nets (1 × 3 m panels) attached to the tethering line of 3 m diameter helium-filled balloons, with each balloon typically carrying three panels. Initially, panels were suspended at 40 m, 120 m, and 160 m agl, but after preliminary results showed higher mosquito panel densities at higher elevations (from August 2013), the typical altitude was 90 m, 120 m, and 190 m agl. When a larger balloon (3.3 m dia.) was deployed at Thierola (August–September 2015), two additional panels were added at 240 m and 290 m agl. Balloons were launched approximately one hour before sunset (~ 17:00) and retrieved one hour after sunrise (~ 07:30), the following morning. To control for insects trapped near the ground as the panels were raised and lowered, comparable control panels were raised up to 40 m agl and immediately retrieved during each balloon launch and retrieval operation. Between September and November 2014, the control panels were raised to 120 m agl. Following panel retrieval, inspection for insects was conducted in a dedicated clean area. Individual insects were removed from the nets with forceps, counted, and stored in labeled vials containing 80% ethanol.

Mosquitoes were morphologically identified to genus, sex, and gonotrophic state before DNA was extracted from whole body of males and separately from abdomen and thorax/head of female mosquitoes. Molecular barcode identification of mosquito species was carried out by PCR amplification of 658bp of the mitochondrial COI gene, amplicons were sequenced, and sequence comparisons of reads with public databases: BOLD and GenBank. Targeted pathogen analysis was performed on head-thorax extractions of females to detect avian and mammalian plasmodia, using 18S RNA pan-*Plasmodium* assay (Mediannikov *et al.*, 2013; Laroche *et al.*, 2017). Sequencing results will be reported elsewhere (Bamou et al.: in preparation).

Data on vectorial status of the mosquito species at altitude was extracted from an extensive literature search summarized by (Lehmann *et al.*, 2021)[preprint]. Here, we included only mosquito species that were reported as probable or likely vectors of pathogens in West Africa based on the references listed in Table S3). For example, our literature records showed that *Ae. mcintoshi* vectors at least 10 MBPs in Africa, but this species was conservatively excluded from the present analysis because no record pertained to West Africa. West Africa was defined to include the following countries: Benin, Burkina Faso, Gambia, Ghana, Guinea, Guinea Bissau, Ivory Coast, Liberia, Mali, Mauritania, Senegal, Sierra Leone, Togo (Lehmann *et al.*, 2021)[preprint]. The evidence used by these sources to implicate vectors includes one or more of the following findings as shown in Table S3: a) multiple isolations of a pathogen from wild vector from whole body or midgut [Win], b) from salivary glands/saliva [WIs], c) from legs [WIL], d) from Thorax and Head [WIth], e) detection of pathogen by circumsporozoite ELISA [WIc], f) laboratory experiments demonstrating vectorial competence (physiological capacity to support the pathogen to the infectious stage and capacity to transmit to a new host, following blood feeding on infectious blood meal), provided the mosquito naturally feeds on relevant hosts [LF], g) transovarial transmission, provided the mosquito naturally feeds on relevant hosts [TF], h) synchronous appearance of vector that feeds on relevant hosts during outbreak/peak transmission [Epi]. Frequently vectors were incriminated by multiple independent studies with either the same or different criteria. The use of “primary” and “secondary” vectors is widespread in the literature, attesting for its utility (Braack *et al.*, 2018; Lehmann *et al.*, 2021; Wilkerson *et al.*, 2021). Despite being vaguely defined, primary vectors are often those that are responsible for the largest share of the transmission (in a particular region). Unless a mosquito species was explicitly indicated as primary vector for that MBP, it was conservatively assumed to be a secondary vector.

### Data analysis

Heterogeneity across species in categorical traits such as sex or exposure to vertebrate blood was evaluated with contingency likelihood ratio chi-square test. If the fraction of cells with expected counts <5 were greater than 20%, we used exact tests based on Monte Carlo simulation of 10,000 samples using Proc Freq (SAS Institute, 2012).

To estimate a species’ mean flight altitude given our panel heights, we used a weighted mean across panel altitudes, whereby each altitude was weighted by the ratio of the proportion of specimens collected at that altitude over the proportion of nights that panel altitude was used throughout the study. The weighed estimate accommodates variation in panel altitude over the course of the study due to systematic changes (i.e., eliminating panel heights of 40 m after the first year of the study) and opportunistic changes (i.e., adding panels at 240 and 280 m when a larger balloon and abundant helium supply were available). If mosquitoes were equally likely to be sampled across all heights, then the ratio would be 1; however, if mosquitoes were more (or less) likely to be caught at particular panel heights then the ratio would deviate accordingly.

To estimate seasonality in high-altitude flight, panel densities of each species (zeros included) were averaged for each month over data across Sahelian stations and years. The log of the monthly mean multiplied by 1,000 (to form integers) was used as a relative index of monthly flight activity accommodating the large variation between species.

The overall risk of a given pathogen to be dispersed at altitude by a mosquito species depends on the abundance of the mosquito at altitude and on its likelihood to carry that pathogen. The logarithm of the total aerial abundance was used as a measure of windborne presence for each mosquito species, and we considered the ratio of 10:1 as the typical difference between primary and secondary vector in their rate of infection with a particular MBPs (see below for sensitivity analysis). The sum of the risk estimates for each pathogen across windborne mosquito vectors (when more than one vector species was indicated for that pathogen) was used to rank pathogens as to their overall risk of windborne spread, and a similar approach was applied to rank the relative contribution of each mosquito species to the overall windborne spread of MBPs. To address the uncertainty and natural variance in differential weights between primary and secondary vectors and assess the sensitivity of the results to error in these weights, we also used estimates that span the range of these weights. We considered a ratio of 4:1 as the minimum differential rate (in rates of infection) in keeping with the definition of primary and secondary vectors. Because typical sample size per species in each study ranges between a few hundreds and a few tens of thousands, the maximum differential rate would be near 10,000:1 (in rates of infection). Because we aim to rank mosquito borne diseases (MBDs) in term of risk for airborne spread by mosquitoes, we applied and compared their ranking based on these three sets weights: 4:1, 10:1, and 10,000:1 (Supp. Results and Discussion) as a sensitivity analysis to address the uncertainty associated with difference between primary and secondary vectors and its effect on our results.

## Results

### The composition of high-altitude mosquitoes in the West African Sahel

A total of 2,340 culicines and 236 anophelines were identified by morphology among the insects collected at altitude during 617 sampling nights on 1,894 sticky nets (panels) in the Sahel of Mali. No mosquitoes were collected on 508 control panels that were raised to 40–100 m agl and immediately retrieved upon launch and retrieval of the standard panels (Huestis et al 2019, Florio et al. 2020), showing that the samples were not contaminated by mosquitoes flying near the ground during the very short duration launching and retrieval procedure. Based on mitochondrial COI sequence similarity, 2,331 specimens (90.5%) were molecularly assigned to seven genera: *Culex* (1,413)*, Aedes* (641), *Anopheles* (236), *Mimomyia* (17), *Lutzia* (13), *Mansonia* (10), and *Eretmapodites* (1), and 2,107 mosquitoes (81.8%) were assigned to 50 species (Table S1). Species represented by a single specimen (N=17, Table S1) probably fly at altitude in lower numbers. To exclude accidental capture, however, we confined our analyses to the species that were represented by at least two specimens in our collection (Fig. 1, Table S1), conservatively assuming that only 33 mosquito species, in six genera are high-altitude migrants. With 106 mosquito species spanning 11 genera reported from Mali (Tandina *et al.*, 2018; Lehmann *et al.*, 2021; Wilkerson *et al.*, 2021), this estimate supports that at least 31% of the Malian mosquito species use this migration modality. Test of homogeneity in this fraction among the three most diverse genera in Mali (N>25) revealed that *Culex* has exhibited higher than expected fraction of species in altitude (52%, P=0.0164, χ^2^_[df=1]_= 5.7, binomial test). *Culex* has also had the largest number of specimens/species (85, Fig. S1). Generally, the mean number of mosquitoes per species at altitude increased across genera with the total number of species found at altitude (r=0.99, P<0.001, N=6).

**Figure 1.**
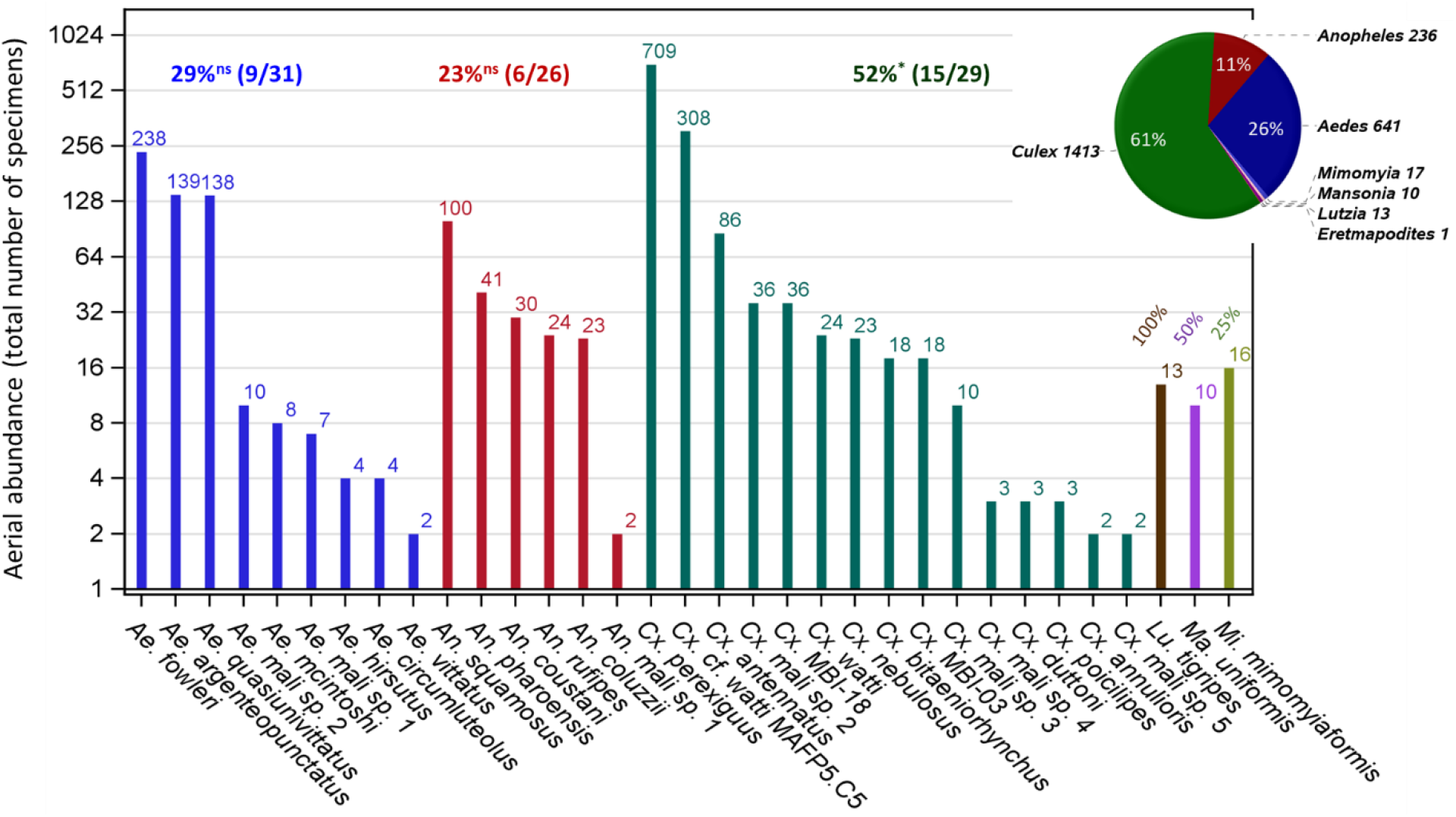
Mosquito composition at altitude (40-290m above ground) in the Mali Sahel (2013–2015) by genera (color) for species represented by two or more specimens in the collection (text). The number of specimens identified using mtCOI DNA barcode similarity is shown above bars (note: logarithmic scale of the Y-axis). The fraction of species found at altitude per genus from the corresponding total known in Mali (see text) are given for each genus above bars and tested against the overall fraction of 28.3% (30/106) using a binomial test (ns and * denote P>0.05 and P<0.05, respectively). The pie chart shows composition of specimens by genera including specimens that were identified by molecular methods (see text).

### Sex ratio and female gonotrophic state composition in high-altitude mosquitoes

Overall, female mosquitoes predominated at altitude across all taxa (85%, N=2,219, Fig. 2), despite significant variation across taxa (P<0.005, Exact χ^2^ test). *Aedes hirsutus* had the lowest female proportion (50%, N=4, Table S2) and four species, including *Ae. mcintoshi and Ma. uniformis* had the highest proportion (100%, N=9, Table S2, Fig. 2). All species with female proportions lower than 70% or higher than 95% had small sample size (4–12, Table S2).

**Figure 2.**
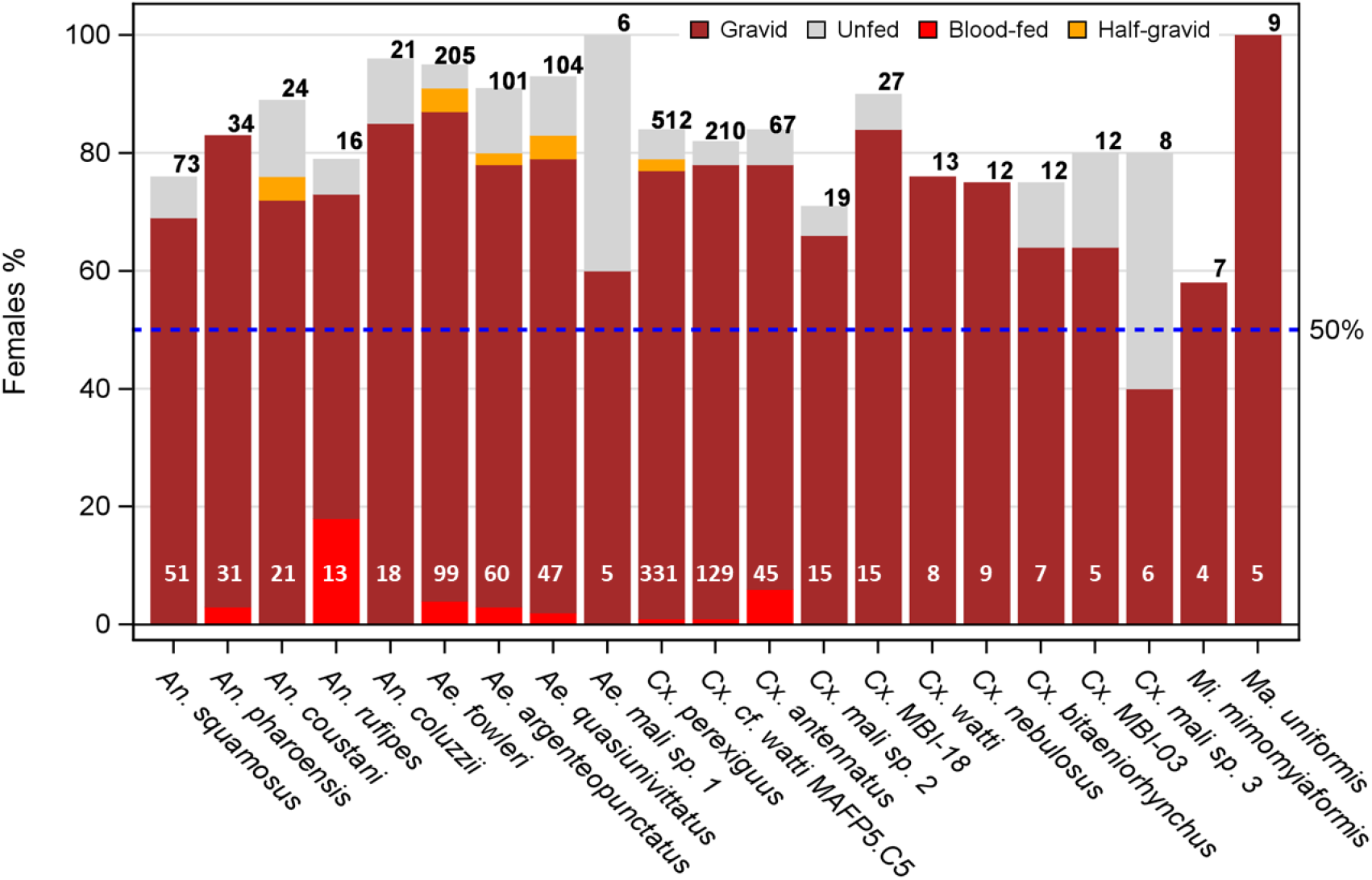
Sex imbalance in mosquitoes at altitude and composition of female’s gonotrophic states by species (note: taxa with <4 females of known gonotrophic state are not shown). The expected equal proportion of females and males is denoted by the broken blue line. Note that except unfed females (gray fill color), all other gonotrophic states must have taken at least one blood meal from a vertebrate host prior to their high-altitude flight and hence could be harboring pathogens from that exposure. Numbers on top (black) and bottom (white) of the bars denote available sample sizes to calculate sex and gonotrophic state composition, respectively.

Gravid females predominated over all other gonotrophic states combined (Fig. 2). Except unfed females, all other gonotrophic states (which must have taken at least one blood meal on a vertebrate host) are hereafter referred to as ‘exposed’. The overall mean proportion of exposed females was 93% (N=924 representing 21 species with sample size of 4 or more females of known gonotrophic state). *Aedes* Mali *sp. 1* had the lowest exposure rate (60%, N=5, Fig. 2), and five species including *An. pharoensis, Cx. watti*, *Cx. nebulosus*, *Mi. mimomyiaformis*, and *Ma. uniformis* had the highest proportion (100%, 4<N<30, Fig. 2). All species with exposure rate below 85% had small sample size (n<10, not shown). However, no heterogeneity was detected in exposure rate among species (P>0.38, Exact χ^2^ test).

### Flight altitude and seasonality of windborne mosquitoes

Mosquitoes were intercepted on all panels set from 40 to 290 m agl. Estimating a species’ typical flight altitude is constrained by the set panel heights (Methods). Thus, although it cannot accurately represent the species actual flight altitude, it can be used to compare species as to their relative typical height weighted by sampling effort at each panel altitude. We estimated the species’ relative mean flight altitude by weighting altitudes based on the ratio of the proportion of mosquitoes captured at that altitude standardized to the proportion of nights that panel altitude was used (Methods, Fig 3a). The weighted mean altitude profile revealed heterogeneity among species as there were non-overlapping 95%CIs. The 95%CI of five species intersected with the mean panel height, but those of three species were below and those of 17 species were above the mean panel height (Fig. 3a, P<0.002, Binomial test of P=0.5). Thus, most culicid species tend to fly at higher altitudes than 150 m (Fig. 3b). Species of *Culex* and *Anopheles* tended to fly higher than species of *Aedes* although the difference was not significant (P=0.071, F_2/17_=3.07 ANOVA excluding genera with 1 species).

**Figure 3.**
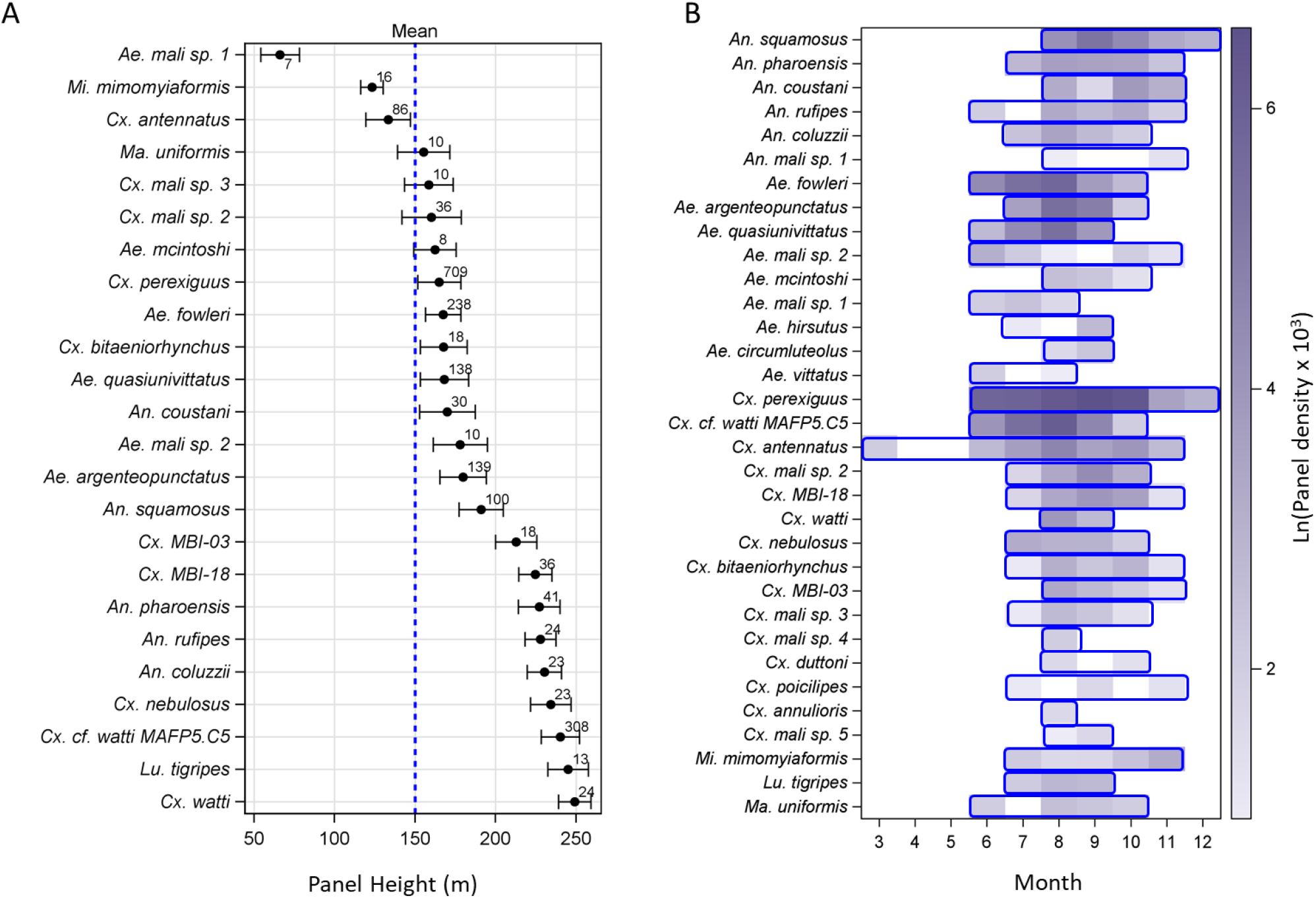
Relative flight altitude (a) and seasonality (b) of high-altitude flying mosquitoes over three years (2013–2015) across the Sahelian aerial stations. Note: species order varies between panels. a) The relative mean flight height of mosquito species (including only those whose N≥5) weighted by the ratio of the proportion of mosquitoes collected on each panel height divided by the proportion of nights this panel height was available and their 95% confidence interval (CI). Overall mean panel altitude is 150 m (denoted by broken blue line). Sample size of each species is given above the mean. b) Seasonality of high-altitude flying Sahelian mosquitoes, showing the period of flight of each species (blue frames), and aerial abundance (filled color intensity), measured as the logarithm of mean panel density * 103 (see text). Note: Sampling was not performed in January–February.

In the Sahel, high-altitude flight was detected across most mosquitoes in June-July and ended October-November (Fig. 3b). Exceptions included *Cx. antennatus* (March–November), *Cx. perexiguus* (June–December) and *An. squamosus* (July–December). Additional sampling would have probably extended these periods, especially for low-abundance species; however, these data suggest a common period of activity across Culicidae (July-October) that coincides with the wet season.

### Assessment of the risk for long-range spread of mosquito-borne pathogens by high-altitude windborne mosquitoes in the West African Sahel

Screening of 921 high-altitude mosquito females using pan-Plasmodium qPCR assay (Laroche et al. 2017), revealed 22 positive samples. Because assays were conducted on thorax-head tissues (Methods), these results indicate the development of the plasmodium to the sporozoite stage, i.e., an overall infectiousness rate of 2.4%. Such results are used to incriminate mosquito vectors of *Plasmodium* spp., rather than a mosquito that was only exposed via infected bloodmeal but cannot transmit it further. Plasmodium-infected high-altitude mosquitoes included members of *Culex* (18), *Aedes* (2), and *Anopheles* (2), with *Cx. perexiguus* comprising ~50% of the infected/infectious mosquitoes. Full detail on pathogen detection in high-altitude mosquitoes will be reported elsewhere (Bamou et al.: in preparation).

To assess the risk for long-range spread of pathogens by the windborne high-altitude mosquito species, we collated literature records (Lehmann *et al.*, 2021) demonstrating involvement in disease transmission by the mosquito species described above. Only species categorized as primary or secondary vectors for a mosquito-borne pathogen (MBP) in West Africa were included (Methods and Table S3). The risk of a given pathogen to be dispersed in altitude by a particular mosquito species depends on the abundance of the mosquito at altitude and on the vector likelihood to carry the pathogen. Accordingly, our risk estimate was approximated by the product of the overall aerial abundance and a weight measuring the relative difference in infectiousness of primary and secondary vectors (1 vs. 0.1, Methods). The sum of the risk estimates for each pathogen across windborne mosquito vectors (when more than one vector species was implicated) was used to rank pathogens by their overall risk of windborne spread, whereas the sum of the risk estimates across pathogens for each mosquito species was used to rank the relative contribution of each mosquito species to the overall windborne spread of mosquito-borne diseases in that region (Fig. 4). To evaluate the sensitivity of the results to different weights, we have compared the rankings of the MBPs and the mosquitoes using differential weights reflecting the range of difference between primary and secondary vectors (Methods and Fig. S2).

**Figure 4.**
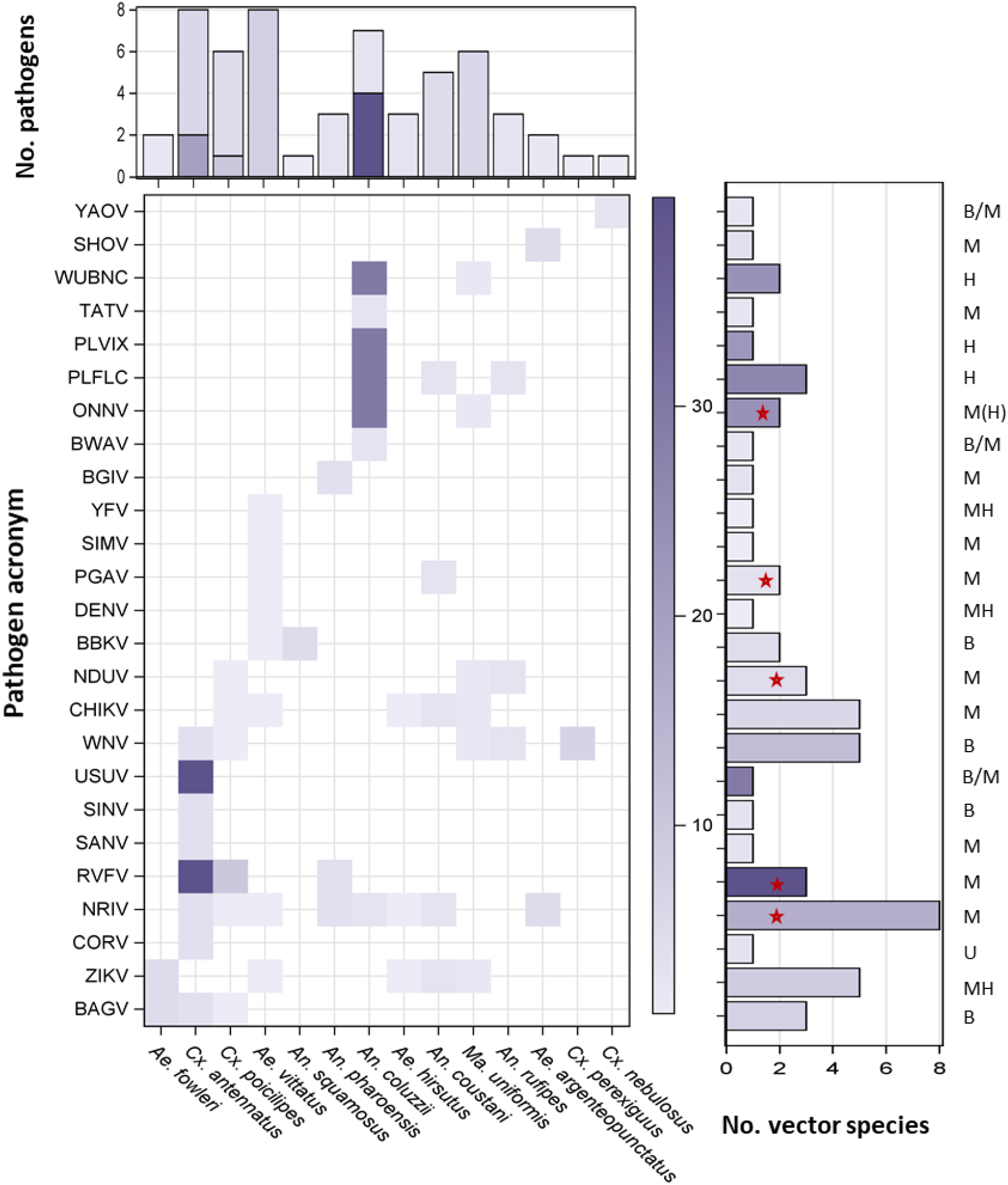
Risk of disease spread by windborne mosquitoes. A heatmap showing pathogen (Y-axis, pathogen acronym, see Table S3) and vector combinations with weight given by the product of vector importance (primary vs. secondary) and aerial abundance (Text). Vertical bars show the number of diseases reportedly transmitted by each vector, either as primary or secondary vector with the total risk weight (color intensity, see text). Horizontal bars show the number of reported vectors of each pathogen, either as primary or secondary vector with the total risk weight (color intensity see text) for each pathogen. Pathogens circulating between humans (H), mammals (M), birds (B), or unknown (U) natural hosts are listed above horizonal bars (M>B denotes mostly mammals but some indication of birds; B/M indicates evidence for both without clear ranking; MH indicate that wild mammals are reservoir hosts but human to human transmission by vector can sustain human infection over several years, M(H) denotes that human-to-human transmission occurs rarely). Red stars indicate MBPs of highest risk for being primarily affected by windborne mosquitoes (see text).

Of the 33 high-altitude flying mosquito species (above), 14 were reported as probable vectors of a total of 25 MBPs in West Africa, including 22 arboviruses, two protozoans (Plasmodia) and one nematode (*Wuchereria bancrofti*, WUBNC, Fig. 4, Table S3). Four mosquito species were recognized as primary vectors for one or more West-African MBPs, namely: *An. coluzzii, An. coustani, Cx. poicilipes*, and *Cx. antennatus*, and the other ten species were categorized as secondary vectors (Fig. 4). Eleven MBPs have been reportedly vectored by one secondary windborne vector species whereas 14 MBPs have been reportedly vectored by multiple vector species or by one primary windborne vector (Fig. 4). The MBPs with elevated risk for windborne spread included all those of highest burden on human health: West Nile virus (WNV), malaria (PLFLC and PLVIX), Zika virus (ZIKV), chikungunya virus (CHIKV), yellow fever virus (YFV), and dengue virus (DENV), as well as Rift Valley Fever virus (RVFV) that threatens domestic animals and humans (Figs. 4 and S2).

That *Ae. vittatus*, *Cx. antennatus*, *Cx. poicilipes, An. coluzzii*, and *Ma. uniformis* are confirmed to undertake long distance migrations is of epidemiological significance in West Africa as these taxa are reported to transmit multiple MBPs (6–8 each), whereas *An. coustani* is reported to transmit five (Fig. 4). *Aedes hirsutus*, *An. pharoensis*, and *An. rufipes* are known to transmit three MBPs each, whereas the remaining six aerial mosquito species contribute only to the transmission of one or two MBPs each (Fig. 4). The index of aerial abundance was not correlated with overall contribution to pathogen spread (r=0.009, P>0.9, N=14).

## Discussion

### Attributes of high-altitude windborne mosquitoes

High-altitude windborne migration of mosquitoes is a dispersal modality of considerable importance in shaping the shifting landscape of mosquito and pathogen composition over space and time, including the generation of disease outbreaks and range expansion (Sellers, 1980; Walsh *et al.*, 1981; Johansen *et al.*, 2003; Sanders *et al.*, 2011; Eagles *et al.*, 2013). It also is bound to affect gene flow between mosquito populations, mediating the spread of insecticide resistance and possibly, in the future, the spread of genetically engineered constructs (e.g., gene drive systems) designed to impact disease control or elimination (Hammond *et al.*, 2016; Epopa *et al.*, 2017; Spitzen & Takken, 2018). Here, we expand our first analysis that covered windborne Anopheline vectors of malaria (Huestis *et al.*, 2019), which consisted of a mere 10% of the mosquitoes in our aerial collection. Leveraging the full collection of mosquitoes that were intercepted at high-altitude (40–290 m agl) over a systematic three-year surveillance period, we provide compelling support for windborne mosquito migration as being i) widespread across diverse genera and species; ii) fivefold more common in females than in males; iii) 13 times more common in females after they took at least one blood meal than in those that had not; iv) typically associated with flight altitudes >150 m and probably exceeding our currently highest trap (290 m); v) coincidental with the wet season (June–November) and peaking in August–October; vi) mediating the spread of plasmodia of vertebrates, and thus representing a proof of principle that pathogens exploit windborne mosquitoes for their own spread, and vii) involving mosquito species that have been reported as vectors to 25 MBPs, including those that inflict the heaviest burden on human and animal health in Africa, such as malaria and Rift Valley fever. Together these findings bolster the role of high-altitude windborne mosquitoes as carriers of pathogens over large distances. Therefore, we propose that aerial sampling, coupled with near real-time holistic pathogen detection approaches, can serve as a powerful monitoring system to map regular pathogen movement, and detect irregular events–functioning as an early warning system to address spillover events, and predict the trajectory of outbreaks and epidemics on regional and continental scales.

The culicid diversity in the aerial collections was high, representing 33 confirmed migrant species and additional 17 likely migrant species of the 106 species reported from Mali (Tandina *et al.*, 2018; Lehmann *et al.*, 2021; Wilkerson *et al.*, 2021) which are equivalent to 31–47% of the country’s fauna. Conservatively, we excluded 17 species represented by a single specimen because some may have been accidental captures (see Suppl. Results and Discussion), however, others were likely less common high-altitude flyers and would be detected through a greater sampling effort. This variation may reflect differences between source population sizes, distances from aerial stations, the fraction of windborne migrants per species, and whether they are diurnal or nocturnal flyers (our sampling was aimed at nocturnal migrants) among other factors. Considering that this three-year collection is only from the Sahel whereas the mosquito fauna of Mali was studied for over a century by many collections using multiple methods targeting diurnal, crepuscular, and nocturnal species from five ecozones, i.e., humid Gallery Forest, wet and dry savannas, Sahel, and desert (Tandina *et al.*, 2018), we expect the actual number of high-flying species in Mali to increase as more studies address this issue. These results suggest that high-altitude flight is a frequent and widespread behavioral trope in mosquitoes, and that this migration modality appears to be part of a mosquito “blueprint”, as also suggested from high-altitude sampling of species of *Culex, Aedes, Anopheles*, and other genera from North America, Asia, Europe, and Australia (Glick, 1939; Reynolds *et al.*, 1996; Kay & Farrow, 2000; Chapman *et al.*, 2004b), and widely known in many groups of insects that track resources in space and time, e.g., locusts, Hemiptera, moths, and other dipterans (Batten, 1967; Riley *et al.*, 1987; Rosenberg & Burt, 1999; Kay & Farrow, 2000; Rose *et al.*, 2000; Holland, 2006; Chapman *et al.*, 2015; Reynolds *et al.*, 2017; Wotton *et al.*, 2019; Florio *et al.*, 2020). It is expected that the abundant seasonal resources available in the Sahel reward long-range migrants, resulting in higher diversity and abundance of migrants (Drake & Gatehouse, 1995; Pedgley *et al.*, 1995; Drake & Reynolds, 2012). Accordingly, the high diversity of insects over the Sahel may reflect migrant mosquitoes from large areas including neighboring ecozones several hundred km away (Garms *et al.*, 1979; Sellers, 1980; Drake & Farrow, 1988; Gatehouse, 1997; Stefanescu *et al.*, 2007; Huestis *et al.*, 2019; Florio *et al.*, 2020). Total aerial abundance varied between culicid species from 2 (e.g., *Ae. vittatus*) to 710 (*Cx. perexiguus*), spanning almost three orders of magnitude (Fig. 1). In each genus the total specimens of 1-2 species outnumbered all remaining specimens of 4–15 species combined. Surprisingly, some of the most common species in ground collections using human and domestic animal baits, or indoor-resting collections in human and animal dwellings (e.g., *Ae. aegypti, Cx. pipiens quinquefasciatus, An. gambiae s.l*.) are less common, rare, or altogether absent in the aerial collection. The abundance of a species at altitude primarily depends on the propensity to engage in high altitude flight and on ground density. If ground densities of these “domesticated species” are among the highest, it seems that their propensity to engage in high altitude flight is relatively low, whereas that of the dominant species (e.g., *Cx. perexiguus*, *Ae. fowleri*, *An. squamosus*) is considerably higher, suggesting that the propensity to migrate varies across mosquito species and that only a fraction of the population engages in windborne migration (partial migration).

Sex ratio at altitude showed that, across species, female mosquitoes outnumber males 5.7 to 1, with modest variation around this ratio (Fig. 2) attributed to small sample size. This ratio likely reflects a fitness differential between migrant female and male mosquitoes. Similarly, a 3:1 ratio of female to male mosquitoes (84:29) was reported in a high-altitude collection from South Asia (Reynolds *et al.*, 1996). A high proportion of females at altitude being gravid (89.6%, Fig. 2) adds weight to the deliberate evolutionary strategy underpinning this long-distance migration. Unlike males, whose fitness depends primarily on the number of virgin females they inseminate, the fitness of gravid females depends primarily on finding larval sites where predation, competition, and pathogen pressures are low and water remains available until larvae complete their development (Service, 1973; Gimnig *et al.*, 2001, 2002; Diabate *et al.*, 2005, 2008; Munga *et al.*, 2006; Mutuku *et al.*, 2006); these are also key drivers of migration in other insects with similar life-histories and habitat requirements (Holland, 2006; Chapman *et al.*, 2012, 2015; Hu *et al.*, 2021). Arrival of migrant females on the winds that also bring the monsoon rains into the Sahel (Maiga *et al.*, 2008; Schneider *et al.*, 2014; Huestis *et al.*, 2019; Florio *et al.*, 2020) ensures finding surface water for oviposition. On the other hand, males’ prospects of mating with a larger number of (virgin) females are considerably lower because typical the sex ratio of offspring is 1:1, suggesting similar competition per female at the destination as around the area of departure. Moreover, the days lost in travel, recovery of energy reserves, and finding mating swarms at destination areas where population density is low probably further depress males’ prospects of mating at its destination, compared with at its site of origin. Thus, the reward of migratory males is the expected larger number of offspring from the fewer successful mates they acquire due to the better conditions in larval sites at the destination. The strong female predominance, therefore, suggests that unless they are propelled to migrate by their pathogens (Hurd, 2003; Altizer *et al.*, 2006; Evans *et al.*, 2009; Martini *et al.*, 2015; Stanczyk *et al.*, 2017; Tallon *et al.*, 2020), the reduction in the number of expected mates of windborne migrating males is large and that the female expected reproductive success is higher, i.e., that most windborne females are capable of at least one successful egg-laying cycle, given their capacity to lay >150 eggs every gonotrophic cycle (Yaro *et al.*, 2006, 2012).

At least 93% of female mosquitoes collected at altitude have taken one or more blood meals on a vertebrate host prior to departure (Fig. 2). The variation associated with this proportion was fully attributed to sample size. Because these diverse mosquito species take blood meals from different domestic and wild mammals and bird species, they are exposed to diverse pathogens found in blood or skin of these diverse hosts. Because of the large daily influx of newly emergent mosquitoes and the time—typically 2–3 days —they take before their first blood meal, this proportion is higher than that found in most ground samples (Gillies & Wilkes, 1965; El-Akad *et al.*, 1988; Lyimo & Takken, 1993; Klowden & Briegel, 1994; Huestis & Lehmann, 2014). The age of these females, however, is important for pathogen detection (Faiman *et al.*, 2020). If the majority are at their first or second gonotrophic cycle, i.e., 2-3 d after their first or second blood meal (~5–6 or 8–9 d old), then pathogen density in infected females would be still low, yielding false negative results for infected mosquitoes that would be found positive a few days later. For example, a typical ELISA or PCR assay using thoracic tissues to detect Plasmodium sporozoites would be negative in females before day 7–9 post infectious bloodmeal (Bass *et al.*, 2008; Hume *et al.*, 2011). Thus, sensitive methods for pathogen detection should be used and pooling mosquitoes should be minimized.

Our results suggest that most mosquitoes tend to fly at higher altitudes than our average panel height (150 m, Fig. 3a). These results agree with radar studies showing that nocturnal insect migrants typically reach heights >200 m above ground and often concentrate in the low-level nocturnal jet that develops above the temperature inversion layer, usually about 200–700 m agl (Reynolds *et al.*, 2005; Aralimarad *et al.*, 2011; Drake & Reynolds, 2012). Therefore, our data underestimate the aerial abundance of most species as well as overall culicid diversity. Species of *Culex* and *Anopheles* appeared to fly higher than species of *Aedes* (P=0.071, F2/18=3.07 ANOVA, excluding genera with 1 species). The variation in typical species flight altitude may be related to the expected displacement due to faster winds at higher altitudes.

As expected for Sahelian insects, high-altitude flight of mosquitoes was markedly seasonal, peaking in August–October and was seldom detected outside June–November (Fig. 3b). The migration period coincides with the availability of surface water for larval sites (above) and with peak disease transmission in the Sahel and surrounding ecozones, thus, increasing the prospects for pathogen transport by windborne mosquitoes. The high-altitude migration period is probably longer than measured in low-abundance species. Indeed, the longest migration period was observed among the most abundant species, e.g., *Cx. perexiguus* (June–December), *Cx. antennatus* (March–November) and *An. squamosus* (July–December). Likewise, the correlation coefficient between the length of the migration period and log of total aerial abundance was 0.59 (N=30, P=0.0006) when the minimum sample size was 2, but fell to 0.4 (N=23, P>0.06) when the minimum total aerial abundance was 10.

### The spread of pathogens by windborne mosquitoes

Finding widespread infection with plasmodial sporozoites (2.4%) in high-altitude mosquitoes across genera (*Culex, Aedes*, and *Anopheles*) is a proof of principle that pathogens of vertebrates are carried by windborne mosquitoes (and presumably by other blood-sucking insects) at altitude. Because finding infected (and infectious) mosquitoes using pan-*Plasmodium* assays and because these assays represent our first targeted screen for pathogens, we expect that additional pathogens including arboviruses and filarial nematodes will also be detected in high-altitude mosquitoes. Full detail on pathogen detection in high-altitude mosquitoes will be reported elsewhere (Bamou et al.: in preparation). Importantly, these results establish that infection with pathogens including the development and amplification of the pathogen in the vector’s body does not prevent mosquito’s high-altitude flight. Nonetheless, comparable data from aerial and ground collections are required to determine whether infection with MBPs suppresses or increases (or does not affect) high-altitude flight in mosquitoes. Unlike laboratory data which remain unresolved (Hurd, 2003; Altizer *et al.*, 2006; Evans *et al.*, 2009; Martini *et al.*, 2015; Stanczyk *et al.*, 2017; Tallon *et al.*, 2020), empirical field data might best address this issue. Because probability of infection increases with age, the age distribution of high-altitude and ground mosquitoes remains key to the interpretation of these infection rates. Finding thoracic infections with plasmodia indicates that at least some of the females are ≥7 days post infectious blood meal and thus are ≥9 days old.

Pathogen transport over tens and hundreds of kilometers in a single night by windborne mosquitoes might result in introduction of MBPs to a vulnerable host population and generate disjoined outbreaks (Garrett-Jones, 1962; Garms *et al.*, 1979; Pedgley, 1982; Service, 1997; Huestis *et al.*, 2019). Such a process of disease spread is considerably faster than by movements of most terrestrial wild hosts and mosquitoes at the ground level. Mixing pathogen genotypes from different localities by windborne transport of MBPs by mosquitoes might also result in new variants. Furthermore, disease control and elimination are complicated by windborne mosquito movement because man-made barriers and boundaries (e.g., insecticide-sprayed zones) are ineffective against windborne mosquitoes. Re-introduction of pathogens into regions after successful local elimination, such as malaria in South Africa may be complicated by windborne mosquitoes (Huestis *et al.*, 2019). Hence, the contribution of windborne plasmodium to the case load of malaria in wild rodents appears small by the number of direct cases compared with local mosquito transmission, but the rare introduction of distinct genotype MBPs from distant sources into other endemic areas can have far reaching implications (Lehmann *et al.*, 2000, 2006; Brasier, 2001; Grünwald *et al.*, 2012; Dardé *et al.*, 2020).

In West Africa, mosquitoes are known to transmit 79 pathogens (Lehmann *et al.*, 2021). Of these, 25 MBPs (32%) are at risk for being transported and introduced into distant sites by one or more of the 14 windborne mosquito species that were reported as probable or partially implicated vectors (Fig 4. and Table S3). Because we conservatively considered only vectors implicated in transmission of MBPs in West Africa that were intercepted at altitude (N≥2) and because the vectors of many MBPs remain unknown, especially those vectors transmitting MBPs among wild vertebrates in sylvatic cycles (Braack *et al.*, 2018; Lehmann *et al.*, 2021; Wilkerson *et al.*, 2021), the actual number of MBPs capable of windborne spread is expected to be higher. Indeed, all the species of plasmodia found to infect our high-altitude mosquitoes represent new vector-MBP records for West Africa (Bamou et al.: in preparation).

This sizable proportion of windborne MBPs and the inclusion of the most burdensome (e.g., YFV, DENV, human malaria) and others posing severe threats for new outbreaks and expansion e.g., RVFV, O’Nyong-Nyong virus (ONNV), highlight the importance of elucidating the impact of windborne mosquito migration on MBD spread and control.

In MBPs circulating between humans as natural hosts i.e., *P. falciparum, P. vivax, W. bancrofti*, and dengue, the importance of spread by windborne mosquitoes is likely smaller than the role of human transport (at least nowadays). Additionally, YFV, ZIKV, CHIKV are maintained among wild mammals (via mosquitoes), but once transmitted to humans, these MBPs can be transmitted between humans (via mosquitoes) for up to several years. Thus, further reducing the number of putative MBPs whose disease ecology is shaped by windborne mosquitoes as a key factor. O’Nyong-Nyong virus is capable of transmission between humans (Rezza *et al.*, 2017), but this appears to be a rare occurrence in relation to the maintenance of the virus in sylvatic cycles, suggesting that the ecology of ONNV is shaped by transmission among nonhuman mammalian hosts. The spread of WNV, which is transmitted among birds, and occasionally transmitted to humans as a zoonosis, is primarily shaped by migratory birds (Sule *et al.*, 2018). However, the case of MBPs that circulate in birds is complex. Firstly, many bird species are residential and move on a distance scale quite similar to resident mammals (Jenkins, 2015; Somveille *et al.*, 2015). Secondly, bird migration often occurs out of synchronization with the transmission season, e.g., the migration from Africa to Europe and Asia occurs in March–May, months before the main transmission season in Sahelian and adjacent ecozones (Jones, 1995). Thirdly, many birds fly several hundreds of kilometers a day and exit the ranges of many MBPs before transmission could take place. Seven MBPs are circulating in birds as natural hosts: BAGV, BBKV, BWAV, USUV, WNV, SINV, YAOV (Fig. 4). Because detailed information on migration of the bird host is lacking, we conservatively excluded all these MBPs from the diseases that are most affected by windborne mosquitoes. The remaining ten MBPs, circulating among wild mammals are less likely to move large distances, and therefore windborne mosquitoes may play a major role in their spatio-temporal dynamics. These include SHOV, NRIV, SIMV, PGAV, TATV, BGIV, RVFV, NDUV, SANV, and ONNV (Fig. 4). Among these ten MBPs, five are transmitted by multiple windborne vectors and thus are ranked as the most affected by this mode of spread: RVFV (with *Cx. antennatus* as primary vector), ONNV (with *An. coluzzii* as primary vector), NRIV (with eight secondary vectors), PGAV, and NDUV (with two and three secondary vectors, respectively; Fig. 4). The long-range spread of RVFV is characterized by disjoined outbreaks (Sindato *et al.*, 2014), as expected when MPB is transmitted by windborne mosquitoes.

The three factors that determine the role of windborne spread of an MBP include: 1) the abundance of the vector species at altitude, 2) the number of windborne mosquito species acting as vectors to this pathogen, and 3) whether the vector is a primary or secondary vector (Fig. 4). Arguably, the epidemiological influence of a single secondary vector might be high even if it introduces a particular MBP into a new host population once every several years, as compared to a single primary vector that does that hundreds of times a year. Fourteen of the 33 high-altitude flying mosquito species identified here, were reported as vectors of one or more MBPs in West Africa (Fig. 4, Table S3). Five mosquitoes —*Ae. vittatus, Cx. antennatus*, *Cx. poicilipes*, *An. coluzzii*, and *Ma. uniformis*—are reportedly vectors for 6–8 MBPs each (Fig. 4), and together contribute to transmission of 22 of the 25 MBPs (Fig. 4), posing a disproportional risk for windborne spread of MBPs. Aside from difference in infection rate, the actual vector status of secondary vectors is typically less established than that of those considered primary vectors. Seventeen of the 25 MBPs are known to be vectored by multiple windborne mosquito species or by a single primary vector (Fig. 4), thus are ranked high in terms of risk of being transported by windborne mosquitoes. They include RVFV, ZIKV, CHIKV, YFV, *P. falciparum, P. vivax*, O’Nyong-Nyong virus (ONNV), Ngari virus (NRIV) whose natural hosts are mammals, as well as WNV and Usutu virus (USUV) whose natural hosts are birds. Many of the MBPs that are transmitted among mammals have been characterized by distant outbreaks – an expected signature of windborne spread, e.g., RVFV, ZIKV, CHIKV, ONNV, NRIV, and YFV, however, this information is not available for many MBPs.

### Conclusion and implications

Despite the low efficiency of our passive aerial sampling method and evidence that mosquito density is likely higher above the height of our highest panel, our results suggest that windborne mosquitoes at altitude represent a key driver of MBP spread at least in the Sahel. Moreover its unique features highlight it as an especially informative monitoring system from public health and food security perspectives because it can reveal the sources and destinations of migrating mosquitoes, pests, and pathogens because of the following reasons: 1) the sizable diversity of mosquito species (and genera) at altitude (which matches or surpasses that of most other sampling methods); 2) the large catchment region from which mosquitoes originate that extends tens and possibly hundreds of kilometers upwind from the sampling station; 3) the predominance of female mosquitoes that have already taken at least one blood meal; 4) the timing of mosquito migration at altitude that coincides with the rainy season when MBP transmission peaks; and 5) the fact that many high-altitude windborne mosquito species (14 of 33) have been fully or partly incriminated as vectors of a total of 25 MBPs in West Africa. These include the MBPs that inflict the heaviest burden on human and animal health (e.g., malaria, yellow fever, dengue, Rift Valley fever), affecting public health and food security. A monitoring system based on high throughput aerial sampling stations will inform about regular mosquito and MBP movement patterns including their major sources, routes, and destinations, which is key to the implementation of successful vector and disease control to reduce buildup of mosquito migrants and the likelihood of carriage of MBPs, as well as prevent arriving immigrants from successful disease transmission and establishment in main destination sites. Moreover, departures from “baseline migration” such as the appearance of a new MBP (or blood/skin pathogen) of vertebrates will be evaluated as a potential new risk, so it could be tracked before it appears in hospitals or veterinary clinics. A prerequisite for such a monitoring system is the demonstration of infection in high-altitude mosquitoes and ideally evidence that it is similar or higher than infection of mosquitoes at ground level. Altogether these results underscore the need to investigate high-altitude windborne mosquitoes and pathogens in Africa to understand the process of disease/outbreak spread, improve disease control strategies, and advance outbreak prediction.

## Acknowledgments

We are grateful to the residents of Thierola, Siguima, Markabougou, and Dallowere for their permission to work near their homes, and for their wonderful assistance and hospitality. We thank to Dr. Moussa Keita, Mr. Boubacar Coulibaly, and Mr. Ousmane Kone for their valuable technical assistance with field and laboratory operations. We thank Drs. Thomas Wellems, Dick Sakai, Laure Juompan, Ms. Fatoumata Bathily, Sekou F Traore, Ms. Margie Sullivan, and Mr. Sam Moretz for vital and often creative administrative and logistic support. This study was supported by the Division of Intramural Research, National Institute of Allergy and Infectious Diseases, National Institutes of Health, Bethesda MD and the Bill & Melinda Gates Foundation (Grand Challenges Explorations grant awarded to TL). Y-ML was supported by the Global Emerging Infections Surveillance Branch of the Armed Forces Health Surveillance Division (AFHSD-GEIS) FY21 project P0030_21_WR and P0065_22_WR. The views, opinions and/or findings contained in this report are those of the author(s) and should not be construed as an official U.S. Department of the Army position, policy, or decision unless so designated by other documentation.

## Supplementary Materials

Diversity, composition, altitude, and seasonality of high-altitude windborne migrating mosquitoes in the Sahel: Implications for disease transmission

### Supplementary Results and Discussion

Because no mosquito was collected on 508 control panels that were raised to 40-100 m agl and immediately retrieved during launch and retrieval of the standard panels, these mosquitoes most probably were captured at altitude rather than near the ground (Huestis et al 2019, Florio et al. 2020). Mosquitoes are not captured on the control panels not only because of their short duration but also because launch stations were set in open fields away from humans, animals, and shelters and because most mosquito species are nocturnal, while launch and retrieval occurred during the day. As it was found in over 55% of the genera, i.e., six (or seven) of the 11 genera in Mali (Lehmann *et al.*, 2021; Wilkerson *et al.*, 2021): *Culex, Aedes, Anopheles, Mansonia, Mimomiya, Lutzia*, and *Eretmapodites* (Table S1), this migration modality is rather common in mosquitoes. Likewise, high proportion of the species in Mali engage in high-altitude migration because depending on whether species sampled by a single specimen are included, our estimate ranges between 31% and 47%. Yet, the actual proportion is expected to be higher because: i) 473 of 2,576 mosquitoes (18.4%) were not assigned to species, and likely include several new species, ii) our aerial sampling was carried out in the Sahel, whereas other ecozones of Mali, which have distinct mosquito fauna have yet to be sampled, and iii) sampling at higher altitudes, e.g., 300-700 m agl, and during stronger winds (precluded given the helium balloons vulnerability to strong winds) would likely increase the number of specimens and the species diversity. The genus Culex predominated in the aerial collection both in terms of the number of mosquito specimens and the number of species. Test of homogeneity among the three largest genera in this fraction revealed that *Culex* has exhibited higher than expected fraction of species in altitude given its total number of species in Mali (45%, P=0.048, η^2^_[df=1]_= 3.9, binomial test). *Culex* has also had the largest number of specimens/species (99).

To ensure we do not include accidentally caught mosquitoes, we excluded species that were represented by a single specimen even though most probably are species that are less abundant in high altitude (see main text). Importantly, two species that appear in our aerial collection as singletons are especially important disease vectors, i.e., *Ae. aegypti, Ae. ochraceus, An. gambiae*, and *Cx. pipiens* (Table S1) (Braack *et al.*, 2018; Lehmann *et al.*, 2021; Wilkerson *et al.*, 2021). An additional concern, we cannot entirely rule out is that during the early phase of the molecular identification, certain specimens used as positive control might have resulted in possible laboratory error due to contamination. Although there is no evidence for this possibility, the inclusion of *Ae. aegypti, An. gambiae*, and *Cx. pipiens* from our laboratory as positive controls requires additional prudence. The *An. gambiae* specimen (Table S1) was separated and identified by another laboratory as previously described (Huestis *et al.*, 2019) prior to the processing of specimens in our own laboratory, precluding this possibility for that species identification.

**Table S1.** Mosquito species in high-altitude in the Sahel and their sample size

| Seq.No. | Genus | Species | No. Specimens |
| --- | --- | --- | --- |
| 1 | *Aedes* | *Ae. fowleri* | 238 |
| 2 | *Aedes* | *Ae. argenteopunctatus* | 139 |
| 3 | *Aedes* | *Ae. quasiunivittatus* | 138 |
| 4 | *Aedes* | *Ae. mali sp. 2* | 10 |
| 5 | *Aedes* | *Ae. mcintoshii* | 8 |
| 6 | *Aedes* | *Ae. mali sp. 1* | 7 |
| 7 | *Aedes* | *Ae. hirsutus* | 4 |
| 8 | *Aedes* | *Ae. circumluteolus* | 4 |
| 9 | *Aedes* | *Ae. vittatus* | 2 |
| 10 | *Aedes* | *Ae. aegypti*<sup>a</sup> | 1 |
| 11 | *Aedes* | *Ae. (Stg.) sp. 1* | 1 |
| 12 | *Aedes* | *Ae. ochraceus* | 1 |
| 13 | *Aedes* | *Ae. triseriatus* | 1 |
| 14 | *Aedes* | *Ae. mali sp. 3* | 1 |
| 15 | *Aedes* | *Ae. mali sp. 4* | 1 |
| 16 | *Aedes* | *Ae. mali sp. 6* | 1 |
| 17 | *Aedes* | *Aedes spp.* | 84 |
| 18 | *Anopheles* | *An. squamosus* | 100 |
| 19 | *Anopheles* | *An. pharoensis* | 41 |
| 20 | *Anopheles* | *An. coustani* | 30 |
| 21 | *Anopheles* | *An. rufipes* | 24 |
| 22 | *Anopheles* | *An. coluzzii* | 23 |
| 23 | *Anopheles* | *An. mali sp. 1* | 2 |
| 24 | *Anopheles* | *An. gambiae* | 1 |
| 25 | *Anopheles* | *An. sp. nr concolor* | 1 |
| 26 | *Anopheles* | *An. mali sp. 2* | 1 |
| 27 | *Anopheles* | *An. cf. coustani 1 NFL-2015* | 1 |
| 28 | *Anopheles* | *Anopheles spp.* | 12 |
| 29 | *Culex* | *Cx. perexiguus* | 709 |
| 30 | *Culex* | *Cx. cf. watti MAFP5.C5* | 308 |
| 31 | *Culex* | *Cx. antennatus* | 86 |
| 32 | *Culex* | *Cx. mali sp. 2* | 36 |
| 33 | *Culex* | *Cx. MBI-18* | 36 |
| 34 | *Culex* | *Cx. watti* | 24 |
| 35 | *Culex* | *Cx. nebulosus* | 23 |
| 36 | *Culex* | *Cx. bitaeniorhynchus* | 18 |
| 37 | *Culex* | *Cx. MBI-03* | 18 |
| 38 | *Culex* | *Cx. mali sp. 3* | 10 |
| 39 | *Culex* | *Cx. mali sp. 4* | 3 |
| 40 | *Culex* | *Cx. duttoni* | 3 |
| 41 | *Culex* | *Cx. poecilipes* | 3 |
| 42 | *Culex* | *Cx. annulioris* | 2 |
| 43 | *Culex* | *Cx. mali sp. 5* | 2 |
| 44 | *Culex* | *Cx. cinereus* | 1 |
| 45 | *Culex* | *Cx. decens* | 1 |
| 46 | *Culex* | *Cx. pipiens<sup>a</sup>* | 1 |
| 47 | *Culex* | *Cx. simpsoni* | 1 |
| 48 | *Culex* | *Culex spp.* | 128 |
| 49 | *Eretmapodites* | *Er. intermedius* | 1 |
| 50 | *Lutzia* | *Lu. tigripes* | 13 |
| 51 | *Mansonia* | *Ma. uniformis* | 10 |
| 52 | *Mimomyia* | *Mi. mimomyiaformis* | 16 |
| 53 | *Mimomyia* | *Mi. mediolineata* | 1 |

<sup>a</sup> It cannot entirely be rule out is that using colony specimens as positive controls may have resulted in erroneous identification. Although there is no evidence for this possibility, additional caution is needed when this species is being considered as a high-altitude migrant.

**Table S2.** Female proportion across species (N≥4) sorted by the proportion of females in the aerial collection.

| Genus | Species | Females | Percent (%) | N |
| --- | --- | --- | --- | --- |
| *Aedes* | *Ae. hirsutus* | 2 | 50.0 | 4 |
| *Lutzia* | *Lu. tigripes* | 5 | 55.6 | 9 |
| *Mimomyia* | *Mi. mimomyiaformis* | 7 | 58.3 | 12 |
| *Aedes* | *Ae. mali* sp. 2 | 5 | 62.5 | 8 |
| *Culex* | *Cx. mali* sp. 2 | 19 | 70.4 | 27 |
| *Culex* | *Cx. nebulosus* | 12 | 75.0 | 16 |
| *Culex* | *Cx. bitaeniorhynchus* | 12 | 75.0 | 16 |
| *Anopheles* | *An. squamosus* | 73 | 76.0 | 96 |
| *Culex* | *Cx. watti* | 13 | 76.5 | 17 |
| *Anopheles* | *An. rufipes* | 16 | 80.0 | 20 |
| *Culex* | *Cx. MBI-03* | 12 | 80.0 | 15 |
| *Culex* | *Cx. mali* sp. 3 | 8 | 80.0 | 10 |
| *Culex* | *Cx. cf. watti* MAFP5.C5 | 210 | 81.7 | 257 |
| *Culex* | *Cx. antennatus* | 67 | 82.7 | 81 |
| *Anopheles* | *An. pharoensis* | 34 | 82.9 | 41 |
| *Culex* | *Cx. perexiguus* | 512 | 83.5 | 613 |
| *Anopheles* | *An. coustani* | 24 | 88.9 | 27 |
| *Culex* | *Cx. MBI-18* | 27 | 90.0 | 30 |
| *Aedes* | *Ae. argenteopunctatus* | 101 | 90.2 | 112 |
| *Aedes* | *Ae. quasiunivittatus* | 104 | 92.9 | 112 |
| *Aedes* | *Ae. fowleri* | 205 | 94.5 | 217 |
| *Anopheles* | *An. coluzzii* | 21 | 95.5 | 22 |
| *Aedes* | *Ae. mcintoshi* | 5 | 100.0 | 5 |
| *Aedes* | *Ae. mali* sp. 1 | 6 | 100.0 | 6 |
| *Aedes* | *Ae. circumluteolus* | 4 | 100.0 | 4 |
| *Mansonia* | *Ma. uniformis* | 9 | 100.0 | 9 |

Sensitivity analysis was used to evaluate the effect of the uncertainty (and natural variance) in the relative difference in likelihood of transmission by primary and secondary vectors on our estimates of importance of windborne spread of different pathogens and on the relative roles of different vectors on overall windborne spread of pathogens (see Main Text). Accordingly, we compared the correlations between our best estimates of windborne spread depicted in Figure 4. with those based on high and low values that span the range of the difference between primary and secondary transmission ratios. We considered an infection ratio (approximately equivalent to transmission contribution ratio) of 4:1 as the minimum differential ratio in keeping with the definition of primary and secondary vectors because similar size difference may also be found between two primary vectors in which one contributes 75% and the other 19%, whereas additional three or more secondary vectors contribute less than the remaining 6%. On the other hand, we consider a ratio of 1:0.0025 among the highest differential rate (in their infection rates) because typical sample size per species in most studies ranges between a few hundreds and a few tens of thousands and the infection rate of the primary vector(s) are near 1% so a ratio of 1:0.0025 implies finding mere 2-3 infected mosquitoes of a sample of 100,000, which very few studies have ever reached not to mention exceeded. Our sensitivity analysis reveals that the estimates of windborne spread based on the mid-range (1:0.1) were highly correlated with those in the extreme low (r=0.99, P<0.0001, N=53) and high of the range (r=0.979, P<0.0001, N=53). When summed over pathogens or mosquito species, high correlations persisted (r>0.95, P<0.0001, N=14, Fig. S1).

**Figure S1.**
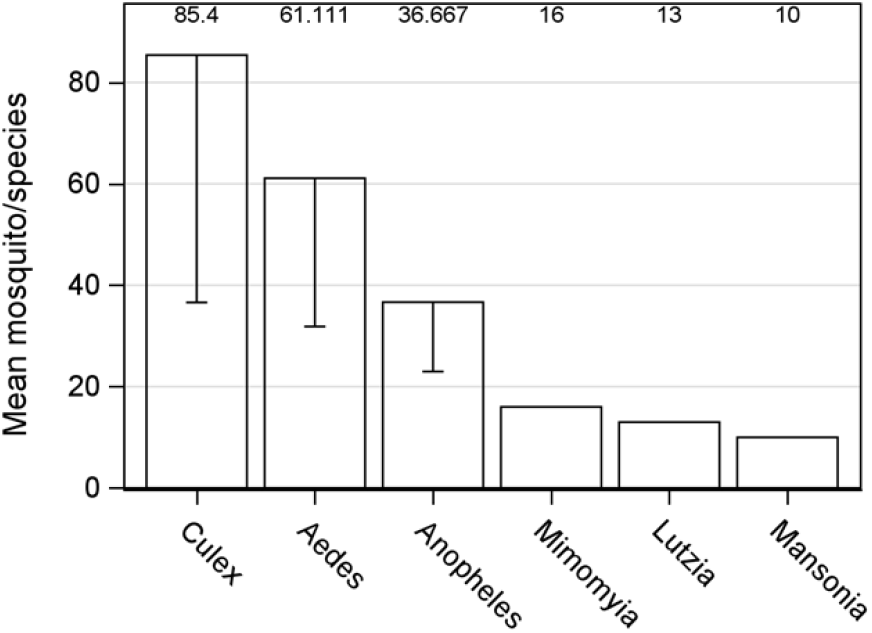
(Supp. Mat.). Mean number of specimen per species across genera and 95% confidence interval for Nspecies/genus > 7. Mean values are shown above bars.

**Figure S2.**
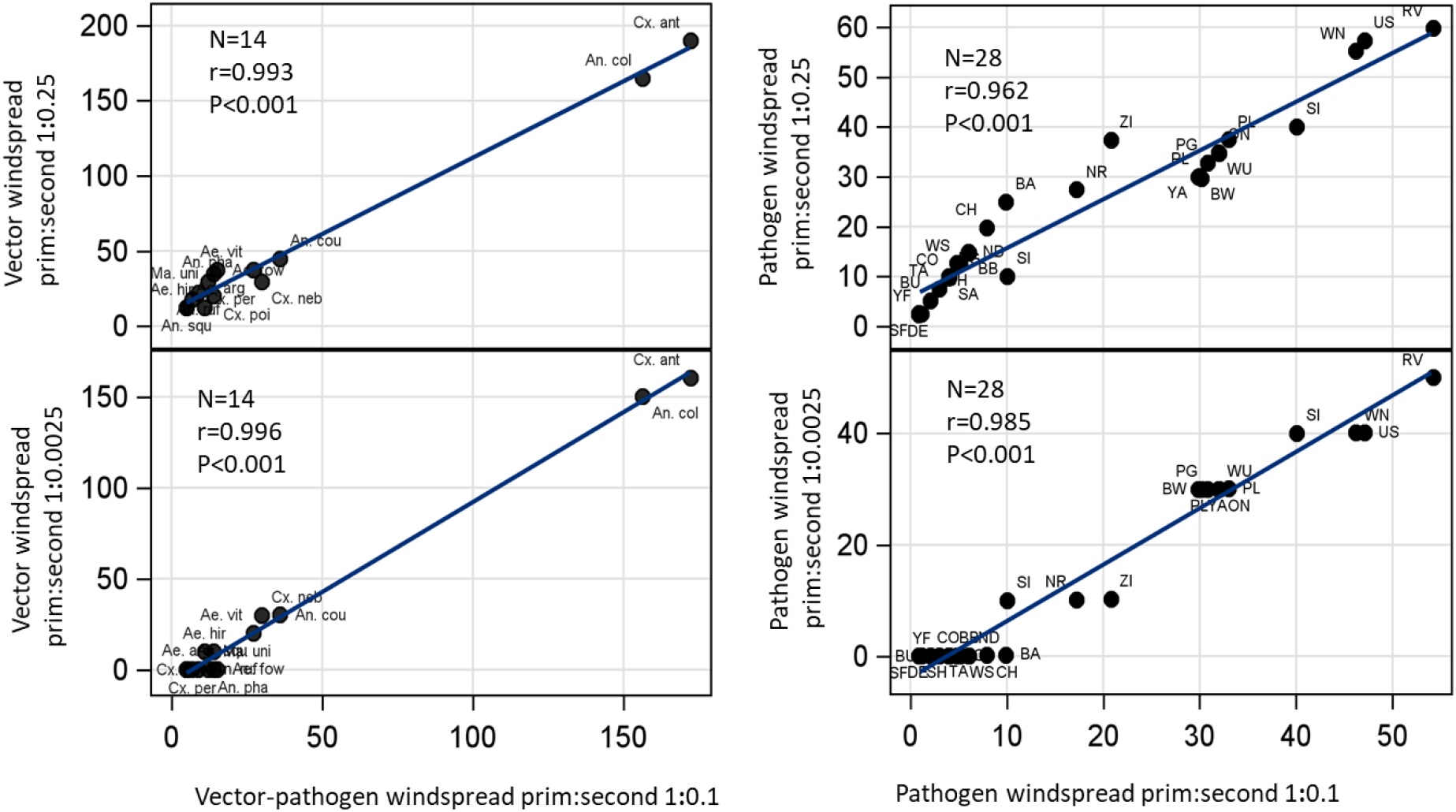
(Also provided as separate figure). Sensitivity of estimates of the relative contribution of windborne transport by different mosquito species (a) and pathogens (b) to the uncertainty in transmission likelihood by primary and secondary vectors (see main text). The relationship between estimates based on the “mid-range” difference between primary and secondary vectors (1:0.1 X-axis), and the lowest difference (1:0.25 Y-axis, top) and highest difference (1:0.0025 Y-axis, bottom) are shown. Values close to the diagonal indicate low sensitivity to different weights or similar patterns across weights. Linear regression between estimators are shown (blue) and observations are labeled by abbreviated vector species and pathogen acronym. Pearson correlation coefficients are shown in top left corner of each panel.

## Literature cited

Altizer, S., Dobson, A., Hosseini, P., Hudson, P., Pascual, M. & Rohani, P. (2006) Seasonality and the dynamics of infectious diseases. Ecology Letters, 9, 467–484.

Aralimarad, P., Reynolds, A.M., Lim, K.S., Reynolds, D.R. & Chapman, J.W. (2011) Flight altitude selection increases orientation performance in high-flying nocturnal insect migrants. Animal Behavior, 82, 1221–1225.

Bass, C., Nikou, D., Blagborough, A.M., Vontas, J., Sinden, R.E., Williamson, M.S., et al. (2008) PCR-based detection of Plasmodium in Anopheles mosquitoes: a comparison of a new high-throughput assay with existing methods. Malaria journal, 7, 177.

Batten, A. (1967) Seasonal movements of swarms of Locusta migratoria migratorioides (R. & F.) in western Africa in 1928 to 1931. Bulletin of Entomological Research, 57, 357.

Braack, L., Gouveia De Almeida, A.P., Cornel, A.J., Swanepoel, R. & Jager, C. De. (2018) Mosquito-borne arboviruses of African origin: Review of key viruses and vectors. Parasites and Vectors, 11, 29.

Brasier, C. (2001) Rapid evolution of introduced plant pathogens via interspecific hybridization. BioScience, 123–133.

Chapman, J.W., Bell, J.R., Burgin, L.E., Reynolds, D.R., Pettersson, L.B., Hill, J.K., et al. (2012) Seasonal migration to high latitudes results in major reproductive benefits in an insect. Proceedings of the National Academy of Sciences, 109, 14924–14929.

Chapman, J.W., Drake, V.A. & Reynolds, D.R. (2011) Recent insights from radar studies of insect flight. Annual Review of Entomology, 56, 337–356.

Chapman, J.W., Reynolds, D.R. & Smith, A.D. (2004a) Migratory and foraging movements in beneficial insects: a review of radar monitoring and tracking methods. International Journal of Pest Management, 50, 225–232.

Chapman, J.W., Reynolds, D.R., Smith, A.D., Riley, J.R., Pedgley, D.E. & Woiwod, I.P. (2002) High-altitude migration of the diamondback moth Plutella xylostella to the UK: a study using radar, aerial netting, and ground trapping. Ecological Entomology, 27, 641–650.

Chapman, J.W., Reynolds, D.R., Smith, A.D., Smith, E.T. & Woiwod, I.P. (2004b) An aerial netting study of insects migrating at high altitude over England. Bulletin of Entomological Research, 94, 123–136.

Chapman, J.W., Reynolds, D.R. & Wilson, K. (2015) Long-range seasonal migration in insects: mechanisms, evolutionary drivers and ecological consequences. Ecology Letters.

Costantini, C., Li, S.G., Torre, A. della, Sagnon, N., Coluzzi, M. & Taylor, C.E. (1996) Density, survival and dispersal of Anopheles gambiae complex mosquitoes in a west African Sudan savanna village. Med. Vet.Entomol., 10, 203–219.

Dao, A., Yaro, A.S., Diallo, M., Timbine, S., Huestis, D.L., Kassogue, Y., et al. (2014) Signatures of aestivation and migration in Sahelian malaria mosquito populations. Nature, 516, 387–390.

Dardé, M.L., Mercier, A., Su, C., Khan, A. & Grigg, M.E. (2020) Molecular epidemiology and population structure of Toxoplasma gondii. Toxoplasma Gondii: The Model Apicomplexan - Perspectives and Methods, 63–116.

Diabate, A., Dabire, R.K., Heidenberger, K., Crawford, J., Lamp, W.O.O., Culler, L.E.E., et al. (2008) Evidence for divergent selection between the molecular forms of Anopheles gambiae: role of predation. BMC Evol Biol, 8, 5.

Diabate, A., Dabire, R.K.K., Kim, E.H.H., Dalton, R., Millogo, N., Baldet, T., et al. (2005) Larval development of the molecular forms of Anopheles gambiae (Diptera: Culicidae) in different habitats: a transplantation experiment. J Med Entomol, 42, 548–553.

Dingle, H. & Drake, A. (2007) What is migration? BioScience, 57, 113–121.

Drake, V.A. & Farrow, R.A. (1988) The influence of atmospheric structure and motions on insect migration. Annual Review of Entomology, 33, 183–210.

Drake, V.A. & Gatehouse, A.G. (1995) Insect migration: tracking resources through space and time. Cambridge University Press, New York.

Drake, V.A. & Reynolds, D.R. (2012) Radar entomology: observing insect flight and migration. CAB International., Wallingford, UK.

Eagles, D., Walker, P.J., Zalucki, M.P. & Durr, P.A. (2013) Modelling spatio-temporal patterns of long-distance Culicoides dispersal into northern Australia. Preventive Veterinary Medicine, 110, 312–322.

El-Akad, a S., Humphreys, J.G. & El-Akad J.G. Humphreys, A.S. (1988) Factors Affecting oviposition and egg production in laboratory-reared Anopheles pharoensis Theobald. Bull Soc Vector Ecol, 13, 243–247.

Epopa, P.S., Millogo, A.A., Collins, C.M., North, A., Tripet, F., Benedict, M.Q., et al. (2017) The use of sequential mark-release-recapture experiments to estimate population size, survival and dispersal of male mosquitoes of the Anopheles gambiae complex in Bana, a west African humid savannah village. Parasites & Vectors, 10, 376.

Evans, O., Caragata, E.P., McMeniman, C.J., Woolfit, M., Green, D.C., Williams, C.R., et al. (2009) Increased locomotor activity and metabolism of Aedes aegypti infected with a lifeshortening strain of Wolbachia pipientis. Journal of Experimental Biology, 212, 1436–1441.

Faiman, R., Yaro, A.S., Diallo, M., Dao, A., Djibril, S., Sanogo, Z.L., et al. (2020) Quantifying flight aptitude variation in wild A. gambiae s.l. in order to identify long-distance migrants. Malaria Journal, 19, DOI 10.1186/s12936-020-03333-2.

Florio, J., Verú, L.M., Dao, A., Yaro, A.S., Diallo, M., Sanogo, Z.L., et al. (2020) Diversity, dynamics, direction, and magnitude of high-altitude migrating insects in the Sahel. Scientific Reports 2020 10:1, 10, 1–14.

Garms, R., Walsh, J.F. & Davies, J.B. (1979) Studies on the reinvasion of the Onchocerciasis Control Programme in the Volta River basin by Simulium damnosum s.l. with emphasis on the sout-western areas. Tropenmed Parasitol, 30, 345–362.

Garrett-Jones, C. (1950) A dispersion of mosquitoes by wind. Nature, 165, 285–285.

Garrett-Jones, C. (1962) The possibility of active long-distance migrations by Anopheles pharoensis Theobald. Bulletin of the World Health Organization, 27, 299–302.

Garrett-Jones, C. & Shidrawi, G.R. (1969) Malaria vectorial capacity of a population of Anopheles gambiae: an exercise in epidemiological entomology. Bull. World Health Organ, 40, 531–545.

Gatehouse, A.G. (1997) Behavior and ecological genetics of wind-borne migration by insects. Annual review of entomology, 42, 475–502.

Gillies, M.T. (1961) Studies on the dispersion and survival of Anopheles gambiae in East Africa, by means of marking and release experiments. Bull. Entomol. Res., 52, 99–127.

Gillies, M.T. & Wilkes, T.J. (1965) A study of the age-composition of populations of Anopheles gambiae Giles and A. funestus Giles in North-Easter Tanzania. BullEntomolRes, 56, 237–262.

Gimnig, J.E., Ombok, M., Kamau, L. & Hawley, W.A. (2001) Characteristics of larval anopheline (Diptera: Culicidae) habitats in Western Kenya. J. Med. Entomol., 38, 282–288.

Gimnig, J.E., Ombok, M., Otieno, S., Kaufman, M.G., Vulule, J.M. & Walker, E.D. (2002) Density-dependent development of Anopheles gambiae (Diptera: Culicidae) larvae in artificial habitats. J Med Entomol, 39, 162–172.

Glick, P.A. (1939) The distribution of insects, spiders, and mites in the air. United States Department of Agriculture, Technical Bulletin. Washington D.C.

Grünwald, N.J., Garbelotto, M., Goss, E.M., Heungens, K. & Prospero, S. (2012) Emergence of the sudden oak death pathogen Phytophthora ramorum. Trends in Microbiology, 20, 131–138.

Hammond, A., Galizi, R., Kyrou, K., Simoni, A., Siniscalchi, C., Katsanos, D., et al. (2016) A CRISPR-Cas9 gene drive system targeting female reproduction in the malaria mosquito vector Anopheles gambiae. Nature Biotechnology 2015 34:1, 34, 78–83.

Harrington, L.C., Scott, T.W., Lerdthusnee, K., Coleman, R.C., Costero, A., Clark, G.G., et al. (2005) Dispersal of the dengue vector Aedes aegypti within and between rural communities. Am J Trop Med Hyg, 72, 209–220.

Holland, R.A. (2006) How and why do insects migrate? Science, 313, 794–796.

Hu, G., Lim, K.S., Horvitz, N., Clark, S.J., Reynolds, D.R., Sapir, N., et al. (2016) Mass seasonal bioflows of high-flying insect migrants. Science, 354, 1584–1587.

Hu, G., Stefanescu, C., Oliver, T.H., Roy, D.B., Brereton, T., Swaay, C. Van, et al. (2021) Environmental drivers of annual population fluctuations in a trans-Saharan insect migrant. Proceedings of the National Academy of Sciences of the United States of America, 118, 2102762118.

Huestis, D.L., Dao, A., Diallo, M., Sanogo, Z.L., Samake, D., Yaro, A.S., et al. (2019) Windborne long-distance migration of malaria mosquitoes in the Sahel. Nature, 574, 404–408.

Huestis, D.L. & Lehmann, T. (2014) Ecophysiology of Anopheles gambiae s.l.: Persistence in the Sahel. Infection, Genetics and Evolution, 28, 648–661.

Huestis, D.L., Yaro, A.S., Traore, A.I., Dieter, K.L., Nwagbara, J.I., Bowie, A.C., et al. (2012) Seasonal variation in metabolic rate, flight activity and body size of Anopheles gambiae in the Sahel. J Exp Biol, 215, 2013–2021.

Hume, J.C.C.C., Hamilton 3rd, H., Lee, K.L.L., Lehmann, T., Hamilton III, H., Lee, K.L.L., et al. (2011) Susceptibility of Anopheles stephensi to Plasmodium gallinaceum: a trait of the mosquito, the parasite, and the environment. PLoS ONE, 6, e20156.

Hurd, H. (2003) Manipulation of Medically Important Insect Vectors by Their Parasites. Annual Review of Entomology, 48, 141–161.

Jenkins, S.H. (2015) Common Patterns in Home Range-Body Size Relationships of Birds and Mammals. https://doi.org/10.1086/283807, 118, 126–128.

Johansen, C.A., Farrow, R.A., Morrisen, A., Foley, P., Bellis, G., Hurk, A.F. Van Den, et al. (2003) Collection of wind-borne haematophagous insects in the Torres Strait, Australia. Medical & Veterinary Entomology, 17, 102–109.

Jones, P.J. (1995) MIGRATION STRATEGIES OF PALEARCTIC PASSERINES IN AFRICA. Israel Journal of Ecology and Evolution, 41, 393–406.

Kay, B.H. & Farrow, R.A. (2000) Mosquito (Diptera: Culicidae) Dispersal: Implications for the Epidemiology of Japanese and Murray Valley Encephalitis Viruses in Australia. Journal of Medical Entomology, 37, 797–801.

Klowden, M.J. & Briegel, H. (1994) Mosquito gonotrophic cycle and multiple feeding potential: contrasts between Anopheles and Aedes (Diptera: Culicidae). J.Med.Entomol., 31, 618–622.

Laroche, M., Almeras, L., Pecchi, E., Bechah, Y., Raoult, D., Viola, A., et al. (2017) MALDI-TOF MS as an innovative tool for detection of Plasmodium parasites in Anopheles mosquitoes. Malaria Journal, 16, 1–10.

Lehmann, T., Blackston, C.R.R., Parmley, S.F.F., Remington, J.S.S. & Dubey, J.P.P. (2000) Strain typing of Toxoplasma gondii: comparison of antigen-coding and housekeeping genes. J.Parasitol., 86, 960–971.

Lehmann, T., Dao, A., Yaro, A.S., Adamou, A., Kassogue, Y., Diallo, M., et al. (2010) Aestivation of the African Malaria Mosquito, Anopheles gambiae in the Sahel. American Journal of Tropical Medicine and Hygiene, 83, 601–606.

Lehmann, T., Kouam, C., Woo, J., Diallo, M., Wilkerson, R. & Linton, Y.-M. (2021) The African mosquito-borne diseasosome: Geographical patterns and range expansion 1. bioRxiv.

Lehmann, T., Marcet, P.L., Graham, D.H., Dahl, E.R. & Dubey, J.P. (2006) Globalization and the population structure of Toxoplasma gondii. Proc Natl Acad Sci U S A, 103, 11423–11428.

Lehmann, T., Weetman, D., Huestis, D.L., Yaro, A.S., Kassogue, Y., Diallo, M., et al. (2017) Tracing the origin of the early wet-season Anopheles coluzzii in the Sahel. Evolutionary Applications, 10, 704–717.

Lyimo, E.O. & Takken, W. (1993) Effects of adult body size on fecundity and the pre-gravid rate of Anopheles gambiae females in Tanzania. Med. Vet.Entomol., 7, 328–332.

Maiga, I.H., Lecoq, M. & Kooyman, C. (2008) Ecology and management of the Senegalese grasshopper Oedaleus senegalensis (Krauss 1877) (Orthoptera: Acrididae) in West Africa. Ann. soc. entomol. Fr. (n.s.), 44, 271–288.

Martini, X., Hoffmann, M., Coy, M.R., Stelinski, L.L. & Pelz-Stelinski, K.S. (2015) Infection of an Insect Vector with a Bacterial Plant Pathogen Increases Its Propensity for Dispersal.

Mediannikov, O., Socolovschi, C., Edouard, S., Fenollar, F., Mouffok, N., Bassene, H., et al. (2013) Common Epidemiology of Rickettsia felis Infection and Malaria, Africa. Emerging Infectious Diseases, 19, 1775.

Miao, J., Wu, Y.-Q., Gong, Z.-J., He, Y.-Z., Duan, Y. & Jiang, Y.-L. (2013) Long-Distance Wind-Borne Dispersal of Sitodiplosis mosellana Géhin (Diptera:Cecidomyiidae) in Northern China. Journal of Insect Behavior, 26, 120–129.

Munga, S., Minakawa, N., Zhou, G., Barrack, O.-O.J.O., Githeko, A.K. & Yan, G. (2006) Effects of larval competitors and predators on oviposition site selection of Anopheles gambiae sensu stricto. J Med Entomol, 43, 221–224.

Mutuku, F.M., Bayoh, M.N., Gimnig, J.E., Vulule, J.M., Kamau, L., Walker, E.D., et al. (2006) Pupal habitat productivity of Anopheles gambiae complex mosquitoes in a rural village in western Kenya. Am J Trop Med Hyg, 74, 54–61.

Pedgley, D.E. (1982) Windborne pests and diseases: Meteorology of airborne organisms. Ellis Horwood Ltd., Chichester.

Pedgley, D.E., Reynolds, D.R. & Tatchell, G.M. (1995) Long-range insect migration in relation to climate and weather: Africa and Europe. In Insect Migration: Tracking resources through space and time (ed. by Drake, V.A. & Gatehouse, A.G.). Cambridge University Press, New York, pp. 3–30.

Reynolds, D.R., Chapman, J.W., Edwards, A.S., Smith, A.D., Wood, C.R., Barlow, J.F., et al. (2005) Radar studies of the vertical distribution of insects migrating over southern Britain: the influence of temperature inversions on nocturnal layer concentrations. Bulletin of Entomological Research, 95, 259–274.

Reynolds, D.R., Chapman, J.W. & Harrington, R. (2006) The migration of insect vectors of plant and animal viruses. Advances in virus research, 67, 453–517.

Reynolds, D.R., Chapman, J.W. & Stewart, A. (2017) Windborne migration of Auchenorrhyncha (Hemiptera) over Britain. European Journal of Entomology, 114, 554–564.

Reynolds, D.R. & Riley, J.R. (1988) A migration of grasshoppers, particularly Diabolocatantops axillaris (Thunberg) (Orthoptera: Acrididae), in the West African Sahel. Bulletin of Entomological Research, 78, 251–271.

Reynolds, D.R., Smith, A.D., Mukhopadhyay, S.S., Chowdhury, A.K., De, B.K., Nath, P.S., et al. (1996) Atmospheric transport of mosquitoes in northeast India. Medical & Veterinary Entomology, 10, 185–186.

Rezza, G., Chen, R. & Weaver, S.C. (2017) O’nyong-nyong fever: a neglected mosquito-borne viral disease, 111, 271–275.

Riley, J.R., Reynolds, D.R. & Farrow, R.A. (1987) The migration of Nilaparvata lugens (stål) (delphacidae) and other hemiptera associated with rice during the dry season in the Philippines: a study using radar, visual observations, aerial netting and ground trapping. Bulletin of Entomological Research, 77, 145.

Rose, D.J.W., Dewhurst, C.F. & Page, W.W. (2000) African Armyworm Handbook: The Status, Biology, Ecology, Epidemiology and Management of Spodoptera exempta (Lepidoptera: Noctuidae). 2nd edn. Natural Resources Institute, University of Greenwich, Chatham.

Rosenberg, J. & Burt, P.J.A. (1999) Windborne displacements of Desert Locusts from Africa to the Caribbean and South America. Aerobiologia, 15, 167–175.

Sanders, C.J., Selby, R., Carpenter, S. & Reynolds, D.R. (2011) High-altitude flight of Culicoides biting midges. The Veterinary record, 169, 208.

Sanogo, Z.L., Yaro, A.S., Dao, A., Diallo, M., Yossi, O., Samaké, D., et al. (2021) The effects of high-altitude windborne migration on survival, oviposition, and blood-feeding of the African malaria mosquito, Anopheles gambiae s.l. (Diptera: Culicidae). Journal of Medical Entomology, 58.

SAS Institute. (2012) SAS software for Windows Version 9.4.

Schneider, T., Bischoff, T. & Haug, G.H. (2014) Migrations and dynamics of the intertropical convergence zone. Nature, 513, 45–53.

Sellers, R.F. (1980) Weather, host and vector--their interplay in the spread of insect-borne animal virus diseases. The Journal of hygiene, 85, 65–102.

Service, M.W. (1973) Mortalities of the larvae of the Anopheles gambiae Giles complex and detection of predators by the precipitin test. Bulletin of Entomological Research.

Service, M.W. (1997) Mosquito (Diptera: Culicidae) dispersal - the long and the short of it. Journal of Medical Entomology, 34, 579–588.

Sindato, C., Karimuribo, E.D., Pfeiffer, D.U., Mboera, L.E.G., Kivaria, F., Dautu, G., et al. (2014) Spatial and Temporal Pattern of Rift Valley Fever Outbreaks in Tanzania; 1930 to 2007. PLOS ONE, 9, e88897.

Somveille, M., Rodrigues, A.S.L. & Manica, A. (2015) Why do birds migrate? A macroecological perspective. Global Ecology and Biogeography, 24, 664–674.

Spitzen, J. & Takken, W. (2018) Keeping track of mosquitoes: a review of tools to track, record and analyse mosquito flight. Parasites & Vectors 2018 11:1, 11, 1–11.

Stanczyk, N.M., Mescher, M.C. & Moraes, C.M. De. (2017) Effects of malaria infection on mosquito olfaction and behavior: extrapolating data to the field. Current Opinion in Insect Science, 20, 7–12.

Stefanescu, C., Alarcón, M. & Àvila, A. (2007) Migration of the painted lady butterfly, Vanessa cardui, to north-eastern Spain is aided by African wind currents. Journal of Animal Ecology, 76, 888–898.

Sule, W.F., Oluwayelu, D.O., Hernández-Triana, L.M., Fooks, A.R., Venter, M. & Johnson, N. (2018) Epidemiology and ecology of West Nile virus in sub-Saharan Africa. Parasites & Vectors 2018 11:1, 11, 1–10.

Takken, W. & Knols, B.G. (1999) Odor-mediated behavior of Afrotropical malaria mosquitoes. Annu.Rev.Entomol., 44, 131–157.

Tallon, A.K., Lorenzo, M.G., Moreira, L.A., Martinez Villegas, L.E., Hill, S.R. & Ignell, R. (2020) Dengue infection modulates locomotion and host seeking in Aedes aegypti. PLOS Neglected Tropical Diseases, 14, e0008531.

Tandina, F., Doumbo, O., Yaro, A.S., Traoré, S.F., Parola, P. & Robert, V. (2018) Mosquitoes (Diptera: Culicidae) and mosquito-borne diseases in Mali, West Africa. Parasites & Vectors, 11, 467.

Touré, H.A., Traoré, K. & Kyei-Baffour, N. (2017) Assessment of changing trends of daily precipitation and temperature extremes in Bamako and Ségou in Mali from 1961-2014. Weather and Climate Extremes, 18, 8–16.

Toure, Y.T., Dolo, G., Petrarca, V., Traore, S.F., Bouare, M., Dao, A., et al. (1998) Mark-release-recapture experiments with Anopheles gambiae s.l. in Banambani Village, Mali, to determine population size and structure. Med. Vet.Entomol., 12, 74–83.

Verdonschot, P.F.M. & Besse-Lototskaya, A.A. (2014) Flight distance of mosquitoes (Culicidae): A metadata analysis to support the management of barrier zones around rewetted and newly constructed wetlands. Limnologica, 45, 69–79.

Walsh, J.F., Davies, J.B. & Garms, R. (1981) Further Studies on the Reinvasion of the Onchocerciasis Control Programme by Simulium damnosum s.I.: The Effects of an Extension of Control Activities into Southern Ivory Coast during 1979, 32, 269–273.

Wilkerson, R.C., Linton, Y.-M. & Strickman, D. (2021) Mosquitoes of the World. Vol. 1 & 2. Johns Hopkins University Press, Baltimore.

Wotton, K.R., Gao, B., Menz, M.H.M., Morris, R.K.A., Ball, S.G., Lim, K.S., et al. (2019) Mass Seasonal Migrations of Hoverflies Provide Extensive Pollination and Crop Protection Services. Current Biology, 29, 2167–2173.e5.

Yaro, A.S., Dao, A., Adamou, A., Crawford, J.E., Traore, S.F., Toure, A.M., et al. (2006) Reproductive output of female Anopheles gambiae (Diptera: Culicidae): comparison of molecular forms. Journal of Medical Entomology, 43, 833–839.

Yaro, A.S., Traore, A.I., Huestis, D.L., Adamou, A., Timbine, S., Kassogue, Y., et al. (2012) Dry season reproductive depression of Anopheles gambiae in the Sahel. Journal of Insect Physiology, 58, 1050–1059.

